# Trazodone rescues dysregulated synaptic and mitochondrial nascent translatomes in prion neurodegeneration

**DOI:** 10.1101/2023.06.05.543732

**Authors:** Hector Albert-Gasco, Heather L. Smith, Beatriz Alvarez-Castelao, Dean Swinden, Mark Halliday, Sudha Janaki-Raman, Adrian Butcher, Giovanna R Mallucci

**Author notes:** Correspondence to: GRM, Full address: Altos Labs, Cambridge Institute, Granta Park, The Portway Building, Great Abington, CB21 6GP, UK.

## Abstract

The unfolded protein response (UPR) is rapidly gaining momentum as a therapeutic target for protein misfolding neurodegenerative diseases, in which its overactivation results in sustained translational repression leading to synapse loss and neurodegeneration. In mouse models of these disorders, from Alzheimer’s to prion disease, modulation of the pathway - including by the licensed drug, trazodone - restores global protein synthesis rates with profound neuroprotective effects. However, the precise nature of the translational impairment, in particular the specific proteins affected in disease, and their response to therapeutic UPR modulation are poorly understood. We used non-canonical amino acid tagging (NCAT) to measure *de novo* protein synthesis in the brains of prion-diseased mice with and without trazodone treatment, in both whole hippocampus and cell-specifically. During disease the predominant translatome changes occur in synaptic, cytoskeletal and mitochondrial proteins in both hippocampal neurons and astrocytes. Remarkably, trazodone treatment for just two weeks largely restored the whole disease translatome in the hippocampus to that of healthy, uninfected mice, predominantly with recovery of proteins involved in synaptic and mitochondrial function. In parallel, trazodone treatment restored the disease-associated decline in synapses and mitochondria and their function to wildtype levels. In conclusion, this study increases our understanding of how translational repression contributes to neurodegeneration through synaptic and mitochondrial toxicity via depletion of key proteins essential for their function. Further, it provides new insights into the neuroprotective mechanisms of trazodone through reversal of this toxicity, relevant for the treatment of neurodegenerative diseases via translational modulation.

## Introduction

Neurodegenerative diseases - including Alzheimer’s, Parkinson’s and related dementias - are characterised not only by the accumulation in the brain of disease-specific misfolded proteins, but also by overactivation of the unfolded protein response (UPR), specifically its PERK (PKR-like endoplasmic reticulum kinase) branch^1–5^. The UPR is a ubiquitous cellular mechanism, its three branches resulting in coordinated signalling cascades aimed at correcting the misfolded protein stress^6, 7^. As part of this response, activation of PERK leads to phosphorylation of its target, eukaryotic initiation factor 2 (eIF2) on its alpha subunit, which blocks translation at the level of initiation reducing global protein synthesis rates in cells. Only a few specific mRNAs escape this translational repression when eIF2α-P levels are high^6, 7^.

In human brains, high levels of PERK-P and eIF2α-P deposition parallel the deposition of disease-associated misfolded proteins^1–5^. While the effects of UPR activation in human disease are unknown, the preclinical data support both a role in pathogenesis and a target for therapy^8–10^. Thus, in animal models of Alzheimer’s disease^11–13^, Parkinson’s^14–16^, ALS^17, 18^, tauopathies^19, 20^ and prion disease^21–25^, chronic PERK pathway activation results in the sustained reduction of global protein synthesis rates associated with subsequent synapse loss and neurodegeneration^14–17, 19, 21–26^. Both genetic^21, 24^ and pharmacological^14–17, 19, 22, 23, 25, 26^ modulation of PERK/eIF2α-P signalling in these mice show cognitive improvement and neuroprotection and recovery of protein synthesis rates in many models^12, 14–17, 20^. In prion disease, this rescues cognitive and synaptic function, prevents synapse loss and neurodegeneration and increases survival^19, 21–25^. Further, in healthy mice, synaptic protein synthesis is essential for learning and memory and reducing eIF2α-P signaling at the synapse boosts cognition in wild type^27–30^ and aged mice^31^. Given the pre-clinical data and the neuropathological findings in human disease, the pathway has become a highly attractive target for the development of new therapies for use across the spectrum of neurodegenerative disease and is a focus for drug discovery^8^. Several compounds, including the small molecule ISRIB^32^ and related compounds restore depressed global translation rates due to high eIF2α-P levels without systemic toxicity in mice and are in development for possible clinical use^33^. Amongst compounds acting in this way is the licensed antidepressant trazodone, which is highly neuroprotective in mouse models of prion disease^22^, frontotemporal dementia^22, 34^ and ALS^18^. Trazodone’s action in UPR/ISR modulation was discovered through unbiased screens^22^, although its precise mechanism of action in reversal of translational repression is not known.

However, despite compelling evidence for the neurotoxic effects of chronic translational repression (which include repression of elongation^35–40^ as well as initiation, amongst other factors) and for the neuroprotective effects of treatments restoring global protein synthesis rates in the brain, the precise nature of the translational decline in disease - and its rescue - are not well understood. Earlier bulk proteomic studies in prion diseased mice showed changes relating to decreased calcium signalling^41^, neuroinflammation and complement activation^42^. Recently, the *de novo* proteomes in mouse models of tauopathy^43^ and the APP/PS1 Alzehimer’s model^44, 45^ (both of which have impaired protein synthesis rates^11, 19, 35, 36^) was analysed, which describes changes in ribosomal^43, 44^ and mitochondrial and synaptic proteins during the course of disease in bulk tissue or hippocampal slices^43–45^. The response of the nascent proteome to modulation of pathways regulating translation in these models was not examined. To better understand the specific dysregulated UPR-related changes in the protein landscape induced during prion neurodegeneration and in response to treatment, we also used non-canonical amino acid tagging (NCAT) to measure *de novo* protein synthesis^46, 47^ in the brains of prion-diseased mice^21^. We analysed changes in the nascent translatome *in vivo,* with and without trazodone treatment during prion-induced neurodegeneration, across the whole hippocampus and also cell-specifically, in neurons and astrocytes respectively. We examined these changes both qualitatively *in situ*, using fluorescence tagging (FUNCAT) and quantitatively, using proteomics after biorthogonal tagging (BONCAT). We found that the decline in protein synthesis during disease reflects, in particular, the significant reduction in many proteins critical for synaptic and mitochondrial function, with an increase in apoptosis signalling in neurons. Critically, these changes are reversed by trazodone treatment, which remarkably largely restores the entire diseased-brain translatomes both in the bulk hippocampal tissue and cell-specifically in hippocampal neurons and astrocytes. In parallel, trazodone treatment had both structural and functional protective effects on synapses and mitochondria, consistent with the changes specific proteins synthesis, restoring reduced numbers of both mitochondria and synapses, and restoring impaired mitochondrial function essential for synaptic and neuronal health. The data bring further granularity and detailed understanding of the proteins affected by translational repression during neurodegeneration across these disorders. Further, for the first time, they show the recovery of the affected proteins and their associated organellar functions in response to therapeutic modulation, supporting the use of drugs such as trazodone to modulate the neurotoxic effects of translational repression.

## Materials and methods

### Prion infection, pharmacological treatment and metabolic labelling

Prion inoculation of tg37^+/-^ mice, NCAT::Camk2a and NCAT::GFAP transgenic mice (Jackson Laboratory, line #028071, #024098, 005359) was performed by inoculating 1% brain homogenate of Chandler/RML prions, as described^48^. Control animals for all strains of mice received 1% normal brain homogenate (NBH) and were culled at the same time points as prion and treated mice. Tg37^+/-^ prion disease mice were dosed intraperitoneally once daily with 40 mg/Kg of trazodone hydrochloride (Merck T6154) or vehicle (saline)^22^ from 8 to 10 weeks post inoculation (w.p.i), and from 16 to 18 w.p.i in NCAT::Camk2a and NCAT::GFAP mice (**Fig. 1A and Supplementary Fig. 1B**), as these mice have a slower prion disease progression. Mice were randomly assigned a treatment by cage number; no mice were excluded from the analysis.

**Figure 1.**
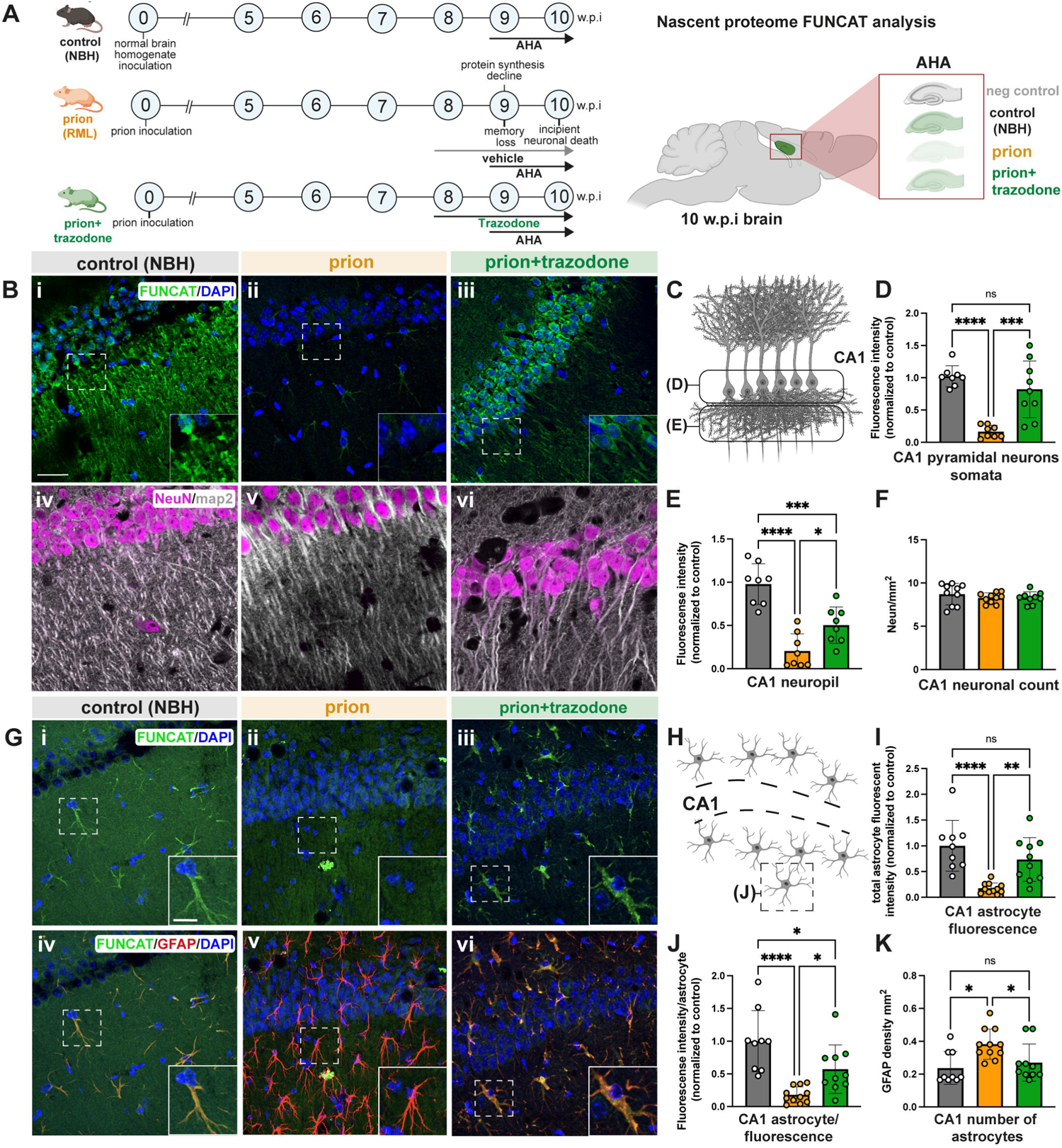
Trazodone treatment restores repressed protein synthesis rates in prion-diseased hippocampal CA1 neurons and astrocytes. **(A)** schematic showing prion disease progression and timing of AHA labelling and trazodone treatment in tg37^+/--^ mice (left). Schematic of sagittal section of the mouse brain (right) showing hippocampal region analysed by FUNCAT. **(B)** Representative images of the nascent proteome labelled by FUNCAT/DAPI (**i-iii**); and NeuN/MAP2-stained dendrites **(iv-vi)** in CA1 pyramidal neurons for all conditions, showing marked reduction during prion disease and restoration by trazodone. **(C)** Schematic of the two regions of CA1 pyramidal neurons analysed. Fluorescence intensity quantification of newly synthesised proteins in CA1 pyramidal neuron somata **(D)** and in neuropil **(E)** for all conditions. **(F)** Neuronal counts in CA1 determined by NeuN staining and quantification. **(G)** Representative images of the nascent proteome in astrocytes labelled by FUNCAT/DAPI (**i-iii**) and merged FUNCAT/GFAP/DAPI signal in control, prion and prion+trazodone **(iv-vi),** showing marked reduction during prion disease and restoration by trazodone. **(H)** Schematic showing location of CA1 astrocytes analysed. **(I)** Total fluorescence intensity for total CA1 astrocytes and **(J)** fluorescence intensity/astrocyte for all conditions. **(K)** Numbers of astrocytes in CA1 region per condition. Scale bar at B(i): 25μm, G(i): inset:10μm., *p* values: * < 0.05, ** <0.001, *** <0.0001,**** <0.00001; ns: non-significant

We used non-canonical metabolic labelling to label the nascent translatome of controls (NBH), prion-diseased (prion) and prion-diseased trazodone treated (prion+trazodone) mice. Methionine surrogates L-azidohomoalanine (AHA) and L-azidonorleucine (ANL) which allow us to label nascent proteins in either non-cell specific or cell-specific manner, respectively, were used. Tg37^+/-^ whole hippocampus non-cell specific nascent translatome labelling was accomplished with 4mM AHA (Fluorochem, # 942518-29-8) mixed with 5% Maltose (Sigma, M9171) in drinking water and in a mash diet from 9 w.p.i to 10 w.p.i. At 10 w.p.i prior to incipient neuronal death^21^ all animals were sacrificed. Cell specific nascent proteome was achieved using a transgene on both NCAT mice containing a point mutated MetRS* at L274G which allows the incorporation of ANL onto met-tRNA (**Supplementary Fig. 1A**). The cell-specific nascent translatome labelling in astrocytes and neurons was achieved by administration of 1% ANL (Iris Biotech, #HAA1625) in drinking water and mash diet^46^. Mice were then culled at 18 w.p.i prior to incipient neuronal death^21^. In parallel all groups had negative ANL/AHA controls to assess methionine surrogate incorporation.

### Fluorescent non-canonical amino-acid tagging (FUNCAT) and immunofluorescence analysis

Neuronal death in tg37^+/-^ mice begins at 10w.p.i, and occurs mainly in the hippocampus, specifically in the CA1 region^21^. Given this, we focused the FUNCAT analysis of the nascent translatome at the CA1 region of the hippocampus.

After metabolic labelling mice were perfused with PBS supplemented with 20mM methionine. Fixed brains were then rinsed in PBS and stored at 4°C until slicing. Brains were mounted on 4% agar, sliced at 40μm using a vibratome (Leica), incubated overnight with 0.5% Triton X100 in PBS, washed extensively with PBS pH 7.8 and clicked overnight with Alexa Fluor™ 647 Alkyne (Thermo Fisher, A10278), triazole ligand (Sigma, # 678937), TCEP (Sigma, #C4706) and copper sulphate (Sigma, #C1297). This was followed by a brief rinse with PBS and mounting with DAPI mounting media. FUNCAT slices were always ‘clicked’ in parallel with negative AHA/ANL control slices. AHA/ANL negative and AHA positive un-clicked slices did not show any FUNCAT signal (**Supplementary Fig. 1D**). After click slices were extensively rinsed and incubated overnight with anti-GFAP (Abcam,#ab4674), anti-MAP2 (Invitrogen, #13-1500) or anti-NeuN (Abcam, ab177487). On the next day slices were incubated with appropriate secondary fluorescent antibodies (Thermo Fisher). After rinsing, slices were mounted on plus-slides (Thermo Fisher), with anti-fade DAPI mounting media Abcam, ab104139) and imaged using a Leica Stellaris SP8 confocal microscope. Z-stack images were taken using a 63x objective with 40 sections at 0.34µM distance/section. Z-stacks were then combined summing fluorescence intensity for each of the sections to a single image.

To measure fluorescence intensity images were analysed with FIJI^49^ by subtracting from the overall intensity the product of the background intensity and total area analysed. CA1 fluorescent intensity at somas was segmented using DAPI and NeuN stain as a reference for hippocampal pyramidal neurons. CA1 fluorescence intensity astrocyte signal was segmented using GFAP astrocyte specific marker as a reference.

### BONCAT and translatome isolation

For BONCAT analysis, ANL and AHA labelled hippocampi were dissected from the rest of the brain. Dissected tissue was then homogenised and lysed in PBS pH 7.4 supplemented with 1% (w/v) Triton-100 and 0.8% (w/v) SDS, protease inhibitors (PI, 1:4000 dilution of protease inhibitor cocktail 3 w/o EDTA, Calbiochem, #539134) and benzonase (1:1000, Sigma) at 75°C for 15min. Lysates were then cleared by centrifugation at RT for 20 minutes and stored at −80°C until further analysis. BONCAT was performed as previously described^46^. In brief lysates were alkylated (iodoacetamide, Sigma, # I1149) for 4h at RT. Following alkylation samples were clean-filtered (PD spintrap G-25 columns, GE Healthcare) and 32μg of protein was clicked to 31μM biotin-alkyne (Jena biosciences, #CLK-TA105) mixed in PBS pH 7.8-PI (Calbiochem, 1:4000), 300μM Triazol ligand (Sigma, #678937) and 84μg/ml CuBr (prepared by a 10mg/ml dilution in DMSO, Sigma, 254185) at RT in the dark overnight.

All biotinylated proteins were then separated using an SDS-PAGE electrophoresis and immunoblotting with anti-biotin (Cell Signalling, #5597) and anti-GAPDH (Santa Cruz Biotechnology, #sc-32233) as loading control. This was followed by the incubation with secondary antibodies Starbright Blue 520 Goat Anti-Mouse (Bio-rad, #12005866) and StarBright Blue 700 Goat Anti-Rabbit (Bio-rad, # 12004161). BONCAT signal for all experimental groups was compared and normalized to their internal AHA/ANL clicked negative controls.

To determine the nature of each of the nascent translatomes, representative mice of each of the AHA/ANL labelled mice cohorts with signal above that of negative controls were chosen for tandem mass spectrometry (LC-MS/MS) analysis. Alkylated and cleaned sample of AHA/ANL incorporated samples were clicked to a disulfide-biotin alkyne (DST) (Click Chemistry Tools, #1498) as described before. Optimal dose of DST alkyne was determined in each case (15µM – 25µM) before upscaling the reactions. Optimal DST alkyne dosages were then used to determine optimal Neutravidin bead (Pierce, #29200) dose for each case (9-15µl dry beads). Upscaling using both optimal dosages of 1200µg of lysate protein was affinity purified for each of the nascent translatomes incubating beads and clicked in PBS pH 7.4 with 0.1% SDS, 1% triton x-100 and PI 1:1000 overnight at 4°C.

Following incubation, beads were extensively rinsed with Neutravidin wash buffer (PBS pH 7.4 1%, triton x-100, 0.2% SDS +PI 1:1000), PBS+PI (1:1000) and 50mM ammonium bicarbonate + PI (1:1000). Following bead rinses, each specific translatome was eluted in 80% volume of dry beads in 5% β-mercaptoethanol, 0.03% SDS in 50mM ammonium bicarbonate +PI (1:1000) mixed in water.

Following elution, quality of the elution was assessed by loading 50µl of the resulting elution on a pre-cast 4-15% gradient gel (Merk, MP41G15) and incubated with SYPRO RUBY (Merck, S4942) protein gel stain ON at RT. On the following day gels were developed according to manufacturer’s instructions (**Supplementary Fig. 2A**). Once the ANL/AHA elutions were seen to be enriched compared to negative controls the rest of the elution was loaded on to gradient gels and sent for LC-MS/MS analysis.

Gel pieces were reduced (DTT) and alkylated (iodoacetamide) and subjected to enzymatic digestion with sequencing grade trypsin (Promega) overnight at 37°C. After digestion, the supernatant was pipetted into a sample vial and loaded onto an autosampler for automated LC-MS/MS analysis.

All LC-MS/MS experiments were performed using a Dionex Ultimate 3000 RSLC nanoUPLC (Thermo Fisher Scientific) system and a Q Exactive Orbitrap mass spectrometer (Thermo Fisher Scientific). Separation of peptides was performed by reverse-phase chromatography at a flow rate of 300 nL/min and a Thermo Scientific reverse-phase nano Easy-spray column (Thermo Scientific PepMap C18, 2mm particle size, 100A pore size, 75 mm i.d. x 50cm length). Peptides were loaded onto a pre-column (Thermo Scientific PepMap 100 C18, 5mm particle size, 100A pore size, 300 mm i.d. x 5mm length) from the Ultimate 3000 autosampler with 0.1% formic acid for 3 minutes at a flow rate of 10 mL/min. After this period, the column valve was switched to allow elution of peptides from the pre-column onto the analytical column. Solvent A was water + 0.1% formic acid and solvent B was 80% acetonitrile, 20% water + 0.1% formic acid. The linear gradient employed was 2-40% solvent B for 30 minutes. Further wash and equilibration steps gave a total run time of 60 minutes.

The LC eluent was sprayed into the mass spectrometer by means of an Easy-Spray source (Thermo Fisher Scientific). All m/z values of eluting ions were measured in an Orbitrap mass analyzer, set at a resolution of 35000 and was scanned between m/z 380-1500. Data dependent scans (Top 20) were employed to automatically isolate and generate fragment ions by higher energy collisional dissociation (HCD, NCE:25%) in the HCD collision cell and measurement of the resulting fragment ions was performed in the Orbitrap analyser, set at a resolution of 17500. Singly charged ions and ions with unassigned charge states were excluded from being selected for MS/MS and a dynamic exclusion window of 20 seconds was employed.

### Analysis of LC-MS/MS data

LC-MS/MS raw data was processed using MaxQuant (Max Planck Institute), matching spectra to the Mus musculus database from Uniprot (Tax ID 10090). Peptide identifications were calculated with FDR<0.01 for multiple comparisons (see further parameters at **Supplementary Table 1**) and proteins with more than 2 peptides were taken further for the analysis. Relative normalized abundance intensity of proteins was given by a label-free quantification approach (LFQ).

Resultant identified proteins peptide intensities were then filtered only for those proteins with higher LFQ values than negative control (**Supplementary Fig. 2B**). Cases in which LFQ values were lower than negative controls were changed to 0. This was followed by a differential expression analysis done using R package DEP^50^ (Bioconductor, 1.19.0). All translatomes were first normalized and imputed (MinProb, q = 0.01). Proteins were considered differentially expressed if α*<0.05*. Hierarchical mappings with dendrograms clustering both LFQ Log 2 normalized intensities and mice as well as PCA plots were generated using DEP package visualization tool. Volcano plots comparing two sets of translatomes were generated using the output DEP results and with Graph-pad prism 9 (all contrasts available in **Supplementary Table 2**).

Following differential protein expression analysis, significantly differentially expressed proteins from all translatomes were analysed for IPA pathway analysis (Qiagen) to highlight activated/inhibited pathways when comparing control (NBH) vs prion, control (NBH) vs prion+trazodone and prion+trazodone vs prion. Representative heatmaps of canonical pathways which were significant and showed predicted activation/inhibition of the pathway were represented using the z-score.

### Immunoblotting

After identification of the different translatomes, validation of protein changes were performed by immunoblotting. Synaptogenesis pathways was validated by loading of 10µg of protein in 4-20% gradient gels from 18w.p.i NCAT::Camk2a mice from each treatment group, and blotted for synaptophysin (Cell signalling, 36406) and Vesicle associated membrane protein 2 (Cell signalling, 13508) levels. GAPDH (Santacruz, sc-32233) was used as a loading control.

Oxidative phosphorylation was validated by loading 10µg of hippocampal protein lysate from 18w.p.i NCAT::Camk2a mice from the three experimental groups and immunoblotting for a OXPHOS mouse cocktail (Abcam, ab110413). Beta actin (Abcam, ab8227) was used as loading control.

### Electron microscopy

Synapse and mitochondrial rescue by trazodone treatment was also validated by scanning electron microscopy (SEM) of CA1 hippocampal pyramidal neurons of control (NBH), prion and prion+trazodone animals.

Prion tg37^+/-^ mice treated with trazodone from 8 to 10w.p.i were compared with prion only and control (NBH) animals (n=3), were transcardially perfused with 2% PFA, 2% glutaraldehyde in 0.9% saline. Perfusions were followed by 24h post-fixation at 4^ο^C and incubated in sodium cacodylate for 4 days. Afterwards, brains were embedded in 4% agar and sliced to 300µM using a vibratome (Leica), osmosized and re-sliced using an ultramicrotome for SEM imaging. The imaging of the CA1 was divided into two regions of interest, first we imaged CA1 at the pyramidal cell layer to include normally degenerating neuronal somas. Second, we imaged synapses at the CA1 stratum radiatum.

Areas of 20mm^2^ were assessed for mitochondrial numbers, and 10mm^2^ for synaptic numbers. Mitochondrial counts have been expressed as the number of mitochondria per pyramidal neurons counted, while synapses counts were expressed per 55µm^2^ to compare with previous studies^26, 51^. Mitochondria were considered present within synapses when found in either a dendrite or an axon at a synapse, expressed per 100µm^2^.

### Mitochondrial isolation and mitochondrial stress test

To assess mitochondrial respiration in prion disease and how it compares to healthy controls and trazodone treated prion-diseased mice, we used mitochondrial isolation combined with a mitochondrial stress test on a Seahorse XF96 Analyzer (Agilent). Isolation of the mitochondria and the mitochondrial stress test followed previously published methods with slight modifications adapted to our specific tissue^52^.

We isolated mitochondria from control (NBH), prion and prion+trazodone hippocampi from mice treated from 8 to 10w.p.i. Each animal was a biological replicate. To isolate hippocampal mitochondria we first dissociated both hippocampi from each animal using mitochondrial isolation buffer (MIB1: 210 mM of d-Mannitol, 70 mM of sucrose, 5 mM of HEPES, 1 mM of EGTA, and 0.5% (w/v) of fatty acid-free BSA pH 7.2) using 15 to 20 stocks with a manual Potter-Elvehjem PTFE pestle (Sigma) using a 8ml glass tube until tissue fully homogenized. This was followed by a series of centrifugations to resolve the mitochondrial fraction from the parts of the cell (800g, 8000g). Once isolated we rinsed the mitochondrial fraction twice with MIB1 and then resuspended in mitochondrial isolation buffer (MAB1: 220 mM of d-Mannitol, 70 mM of sucrose, 10 mM of KH2PO4, 5 mM of MgCl2, 2 mM of HEPES, 1 mM of EGTA, and 0.2% (w/v) of fatty acid-free BSA, pH: 7.2). Once ready for testing we quantified the amount of mitochondria from each isolation using Bio-Rad Protein Assay Kit (Bio-Rad, #5000001), plated 3µg/50µl/well onto a 96 well plate with all animals having 6-8 technical replicates (101085-004, Agilent). Mitochondrial-plates were then centrifuged at 2000g, 4°C, for 20 minutes. After centrifugation we incubated the mitochondrial-plate in a non-CO2 incubator at 37°C for 10 minutes before running the mitochondrial stress test.

Mitochondrial response to the differing stressors was assessed using an established protocol^52^. In brief, we first measured basal respiration rate in the presence of all substrates (10mM succinate, 5mM malate and 10mM glutamate), followed by injection of 4mM ADP (Sigma, A2754), 5µM Oligomycin (Sigma, O4876), 5µM FCCP (Sigma, C2920) and 5µM Rotenone + Antimycin A (Sigma, R8875, A8674). As a quality control of the isolated mitochondria we stained with 200µM MitoTracker® Red CMXRos mixed in MAB1 (Cell signalling, #9082) showing intact mitochondria for all conditions (**Supplementary Fig. 4D**). Basal respiration was calculated by subtracting basal OCR minus the OCR after injection of rotenone and antimycin A. Respiration after ADP injection was calculated as the subtraction of ADP OCR minus the OCR after injection of rotenone and antimycin A. Proton leak respiration was calculated as the OCR after oligomycin injection minus the OCR after injection of rotenone + antimycin A. Maximal respiratory capacity was measured as the OCR value after FCCP injection minus the OCR after rotenone + antimycin A injection.

### Statistical analysis

Statistical analysis of FUNCAT and BONCAT analysis to assess AHA/ANL incorporation was done using a one-way ANOVA followed by a tukey multiple comparison test with an alpha of 0.05, analysed using Graph pad prism 9. Differential expression analysis on nascent translatomes was done using Stats and limma (Bioconductor 3.17), with R packages contained within the DEP package^50^. Statistical analysis of significantly enriched pathways (p<0.05) from significant proteins was obtained from ingenuity pathway analysis (IPA). Significant analysis of interacting clusters was obtained through metascape-String analysis^53^. Statistical analysis of validation immunoblots for synaptic proteins and OXPHOS components were done using a one-way ANOVA followed by a tukey multiple comparison test with an alpha of 0.05 analysed using Graph pad prism 9. SEM differences for both synapse and mitochondrial counts were done using a one way ANOVA followed by a tukey multiple comparison test with an alpha of 0.05, using Graph pad prism 9. Statistical differences for mitochondrial respiration were assessed with One way ANOVA followed by a tukey multiple comparison test with an alpha of 0.05 analysed using Graph pad prism 9.

## Results

### FUNCAT analysis shows trazodone restores repressed global protein synthesis rates in prion-diseased CA1 neurons and astrocytes

We used tg37^+/-^ mice^54^ inoculated with Rocky Mountain Laboratory (RML) prions for whole hippocampal analyses (see schematic, **Fig. 1A**). These transgenic mice, used extensively in our previous studies^21–25, 48, 51, 54^, over-express prion protein (PrP) ∼3 fold and have a relatively rapid disease course, dying at 12 weeks post inoculation (w.p.i.). They show onset of UPR activation and protein synthesis rates decline from 9 w.p.i, along with cognitive impairment and behavioural change and synaptic dysfunction, with onset of neuronal death in the hippocampal CA1-3 region beginning at 10 w.p.i., followed by overt neurodegeneration and clinical signs of prion disease and death at 11-12 w.p.i^21^. We used NCAT to determine the changes in the nascent translatome in whole tissue due to UPR activation, and its response to treatment to better understand the specific processes leading to neuronal death in prion neurodegeneration. Thus, the methionine analogue, L-azidohomoalanine (AHA), when incorporated into proteins allows either fluorescence or biotin labelling of AHA-containing proteins through click chemistry. Fluorescence-AHA-labelled proteins can be visualised using FUNCAT, while biotin-AHA labelled proteins can be affinity purified and further analysed using BONCAT^46, 47^ (see schematic in **Supplementary Fig. 3A**).

We first visualised global protein synthesis rates, using FUNCAT, in hippocampal sections from prion-infected tg37^+/-^ mice treated with vehicle or trazodone from 8 w.p.i., and a control group inoculated with normal brain homogenate (NBH). AHA was administered between 9 and 10 w.p.i. in all groups to ensure widespread incorporation during the period of UPR activation and translational repression that leads to incipient neuronal loss (**Fig. 1A**). Animals were sacrificed at 10 w.p.i. for analysis and their hippocampi examined. We found a dramatic decline in global protein synthesis rates in the CA1 region pyramidal neuron cell bodies of prion-diseased mice, with average fluorescence intensity (FI) of 0.17 compared to FI of 1.02 in uninfected controls (*p* <0.0001) (**Fig. 1B, panels i, ii and Fig. 1D).** This decline was also seen in neurites: FI 0.2 in prion versus FI 0.98 in controls (*p* <0.0001) (**Fig. 1B, panels i, ii**). Remarkably, treatment with trazodone restored repressed protein synthesis rates almost completely in neuronal cell bodies: FI 0.82 vs 0.17 untreated (*p =* 0.0003) and partially in the neuropil: FI 0.5 vs 0.17 in untreated; *p* = 0.0301) (**Fig. 1B, panels i-iii and Fig. 1D,E**). The reduction - and restoration - of protein synthesis rates was not due to changes in neurite density in the different conditions, as confirmed by comparable levels of NeuN/MAP2 labelling in neurites in all experimental groups (**Fig. 1B, panels iv-vi**), nor to changes in CA1 pyramidal neuron density (**Fig. 1F**).

Astrocytes are also affected by prion disease, showing marked proliferation and activation^24, 48, 55^ (**Fig. 1G, I-K**). Indeed, astrocytic UPR activation leads to a reactive state that contributes to neurodegeneration via reduced trophic support to neurons via altered secretome resulting from reduced translation rates^24^. Co-labelling with FUNCAT and the astrocytic marker, GFAP, in CA1 sections revealed marked repression of the nascent translatome in astrocytes in prion-disease, consistent with our previous findings^24^, which was restored by treatment with trazodone (**Fig. 1G, panels i–vi**). Fluorescent intensity quantification (**Fig. 1H**) confirmed a lowered rate of nascent protein synthesis in prion disease vs control, both per astrocyte: FI 0.18 vs 1.00 (*p* <0.0001) and for all astrocytes: FI 0.17 vs 1.00; (*p* <0.0001), respectively **(Fig. 1I-J),** which was largely restored by trazodone treatment: FI 0.75/astrocyte vs FI 1.00 in controls (*p* = 0.0352) and FI 0.73 for all astrocytes vs FI 1.00 in controls; *p =* 0.005) **(Fig. 1I-J).** In untreated mice, the decline of the FI signal occurred despite a significant increase in density of astrocytes, consistent with reduced translation rates in activated diseased astrocytes (**Fig. 1K)**, again in agreement with previous findings^24^.

In parallel, we performed FUNCAT on mice in which we labelled the nascent proteome in a cell-specific manner. We used NCAT-mice^47^, which, which when crossed with specific Cre-expressing lines, in this case CaMK2a-Cre and GFAP-Cre, express mutated methionine tRNA synthetase, MetRS*, in forebrain neurons and astrocytes, respectively (**Supplementary Fig. 1A)**. This protein will only incorporate the methionine analogue, azidonorleucine (ANL), into new proteins, similar to AHA. NCAT::CamK2a and NCAT::GFAP mice were inoculated with RML prions and treated either with trazodone or vehicle, as before. Controls were inoculated with NBH (**Supplementary Fig. 1B**). NCAT and the Cre lines do not over-express PrP and thus have a slower incubation period than tg37^+/-^ mice, typical for wild type C57/Bl6 mice. Accordingly, UPR activation and translational repression occurs at 17 w.p.i., with neuronal loss beginning around 18 w.p.i. and terminal clinical signs at 22 w.p.i^21^. Consistent with this, FUNCAT-labelling with ANL of the CaMK2a-specific nascent translatome from 17 w.p.i. to 18 w.p.i. confirmed marked repression of protein synthesis rates in CA1 pyramidal neurons, which was restored by trazodone treatment (**Supplementary Fig. 1B, panels i-iii**), equivalent to the changes seen in tg37^+/-^ mice at 9w.p.i to 10 w.p.i. (**Fig. 1B, panels i-iii)**. Similarly, FUNCAT-labelling with ANL of the astrocyte-specific nascent translatome showed repression in global protein synthesis rates at 17 w.p.i. in prion-diseased CA1 astrocytes and its restoration with trazodone treatment at 18 w.p.i. (**Supplementary Fig. 1C, panels iv -vi**). (Background fluorescent signal due to ‘click’ effect was negligible for all experimental groups (**Supplementary Fig. 1D, panels i-vi**)).

### BONCAT analysis shows trazodone restores the repressed nascent proteomes of neurons, astrocytes and whole hippocampus in prion-disease

We next used BONCAT to isolate the individual nascent proteomes followed by LC-MS/MS from AHA-labelled whole hippocampus (tg37^+/-^ mice) and ANL-labelled hippocampal neurons (NCAT::CaMK2a mice) and astrocytes (NCAT::GFAP mice) for each of the three experimental conditions (NBH control, prion-diseased, prion-disease+trazodone treatment) (see schematic, **Fig. 2A**). All nascent translatomes had an average signal above background and confirmed findings from FUNCAT analysis (**Supplementary Fig. 2)**. We identified a total of 1100 proteins in the nascent translatomes of whole hippocampi from tg37^+/-^ mice, of which 716 were in control samples, 982 in prion-only and 949 in prion+trazodone samples. In the neuron-specific nascent translatomes, we identified 830 proteins of which 700 were in controls, 781 in prion-only and 785 in prion+trazodone treated mice. In the astrocyte-specific nascent translatomes there were 924 proteins of which 746 proteins were identified in controls, 745 proteins in prion-only and 814 in prion+trazodone samples (**Fig. 2B;** full list in **Supplementary Table 2**). BONCAT quality control on elutions was confirmed by Sypro Ruby protein staining and log2 intensity enrichment plots (**Supplementary Fig. 2A and B).** Further, the neuronal and astrocytic nascent translatomes were positive for their corresponding cell-specific markers (**Supplementary Fig. 2C**), and the whole hippocampal translatome included markers of oligodendrocytes, microglia and endothelial cells in the (see **Supplementary Table 2)**.

**Figure 2.**
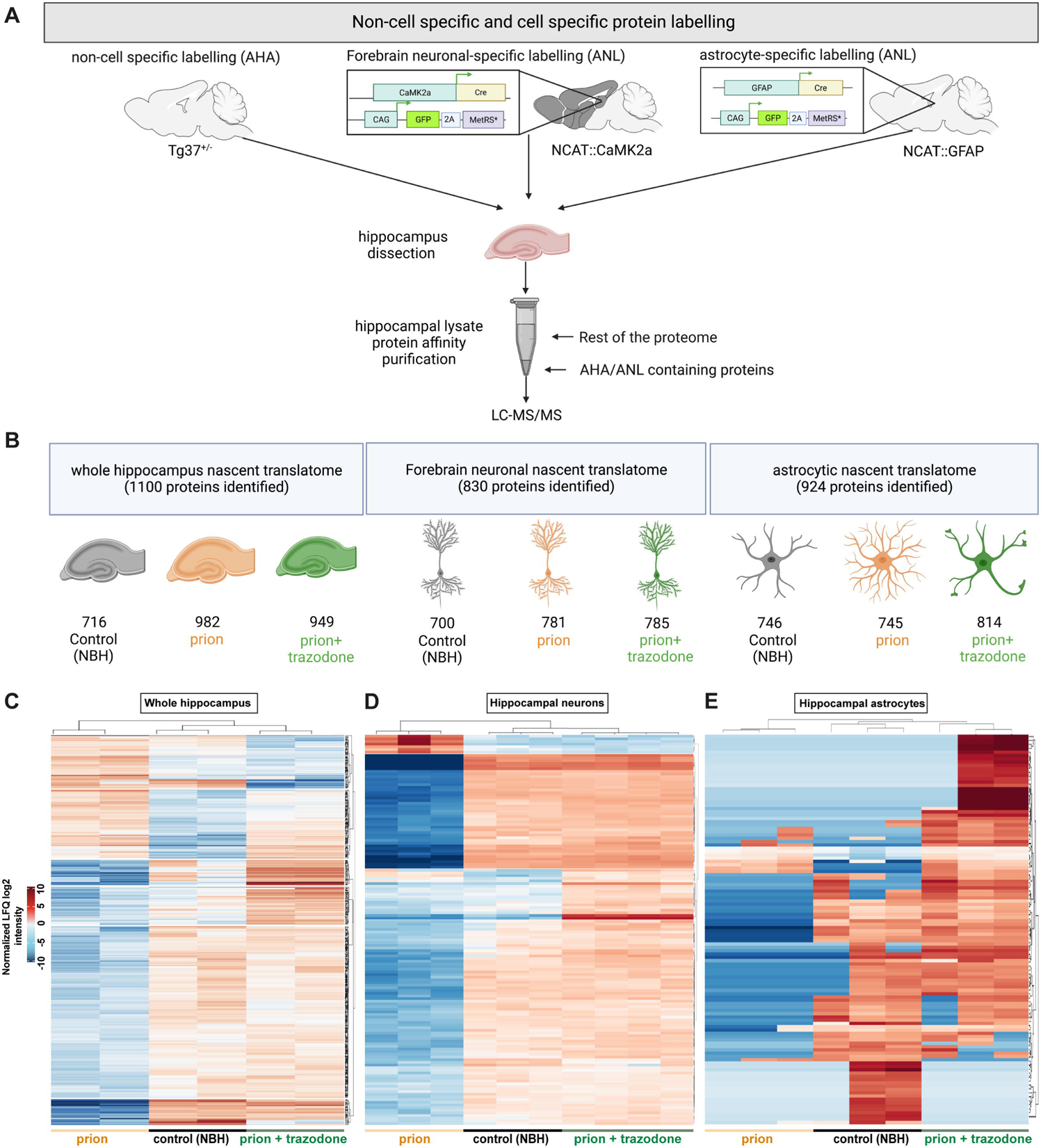
Trazodone restores the nascent proteomes of neurons, astrocytes and whole hippocampus in prion-diseased brains. **(A)** Schematic showing experimental steps for BONCAT analysis: AHA/ANL labelling of the mouse brain, hippocampal dissection, followed by affinity purification of AHA/ANL labelled proteins. **(B**) Total numbers of proteins detected in control (NBH), prion and prion+trazodone treatment groups by LC-MS/MS in the whole hippocampus, in neurons and in astrocytes, respectively. **(C)** hierarchical heatmap by log2 normalized LFQ intensity from whole hippocampus, (**D**) neurons and (**E**) astrocytes, showing clustering of proteins and mice per experimental group. Differentially expressed significant proteins: *p* value < 0.05.

We used label free quantification (LFQ) values to analyse relative differences in abundance of proteins between the different nascent translatomes. When represented in hierarchical heatmaps, LFQ normalized values of significantly expressed proteins showed clustering of values from control, prion and prion+trazodone animals for each condition (**Fig. 2 C,D,E**). Prion disease caused overall reduction (blue cells) in many proteins across each condition compared to high levels (red cells) in control animals. Heatmaps from trazodone-treated animals were very similar to those of controls across each condition, with the majority of proteins restored to control levels, other than a small proportion (see **Supplementary Table 2** and **Figs. 3** and **4** for details). The astrocytic translatome showed more variability in disease versus control and trazodone-treated mice, reflecting the phenotypic variability and mixed profiles derived from varying activation states of astrocytes in disease^56, 57^, but the groups still cluster as the others **(Fig. 2D)**. In line with these data, analysis of variance by principal components (PCA), control and prion+trazodone cluster separately to the translatomes of untreated prion-diseased animals for both the whole hippocampus and hippocampal neurons, but are less clear-cut for astrocytes (**Supplementary Fig. 3D)**.

**Figure 3.**
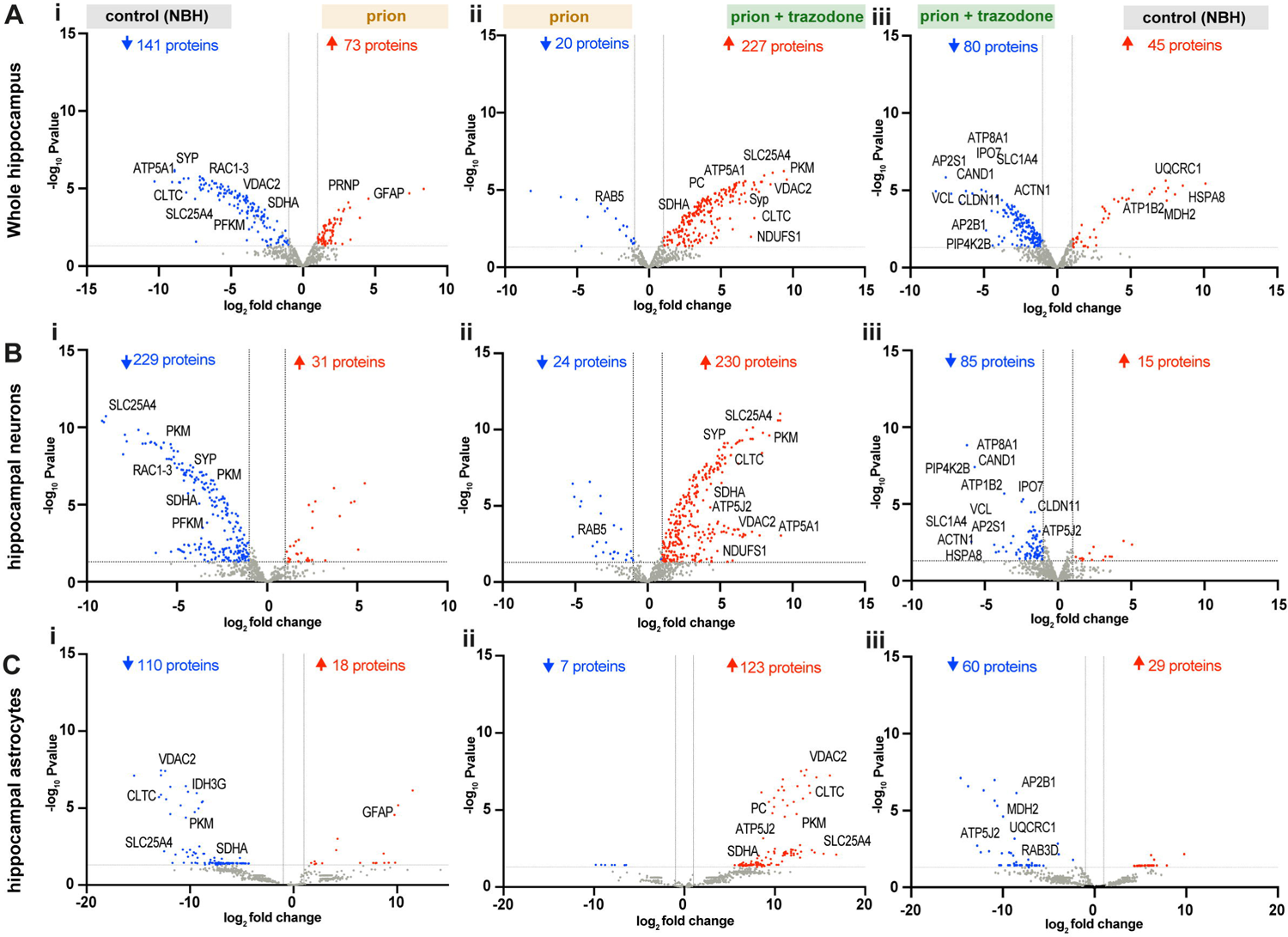
Trazodone treatment restores specific subsets of proteins in the whole hippocampus and cell-specifically, in neurons, astrocytes. **(A)** volcano plots of whole hippocampus translatomes comparing (**i**) prion with control (NBH) translatomes, (**ii**) prion+trazodone with prion and (**iii**) control (NBH) with prion+trazodone. **(B)** volcano plots of hippocampal forebrain neuronal translatomes comparing (**i**) prion with control (NBH) translatomes, (**ii**) prion+trazodone with prion and (**iii**) control (NBH) with prion+trazodone. **(C)** volcano plots of hippocampal astrocytic translatomes comparing prion with (**i**) control (NBH) translatomes, (**ii**) prion+trazodone with prion and (**iii**) control (NBH) with prion+trazodone. Differentially expressed significant proteins: *p* value < 0.05.

**Figure 4.**
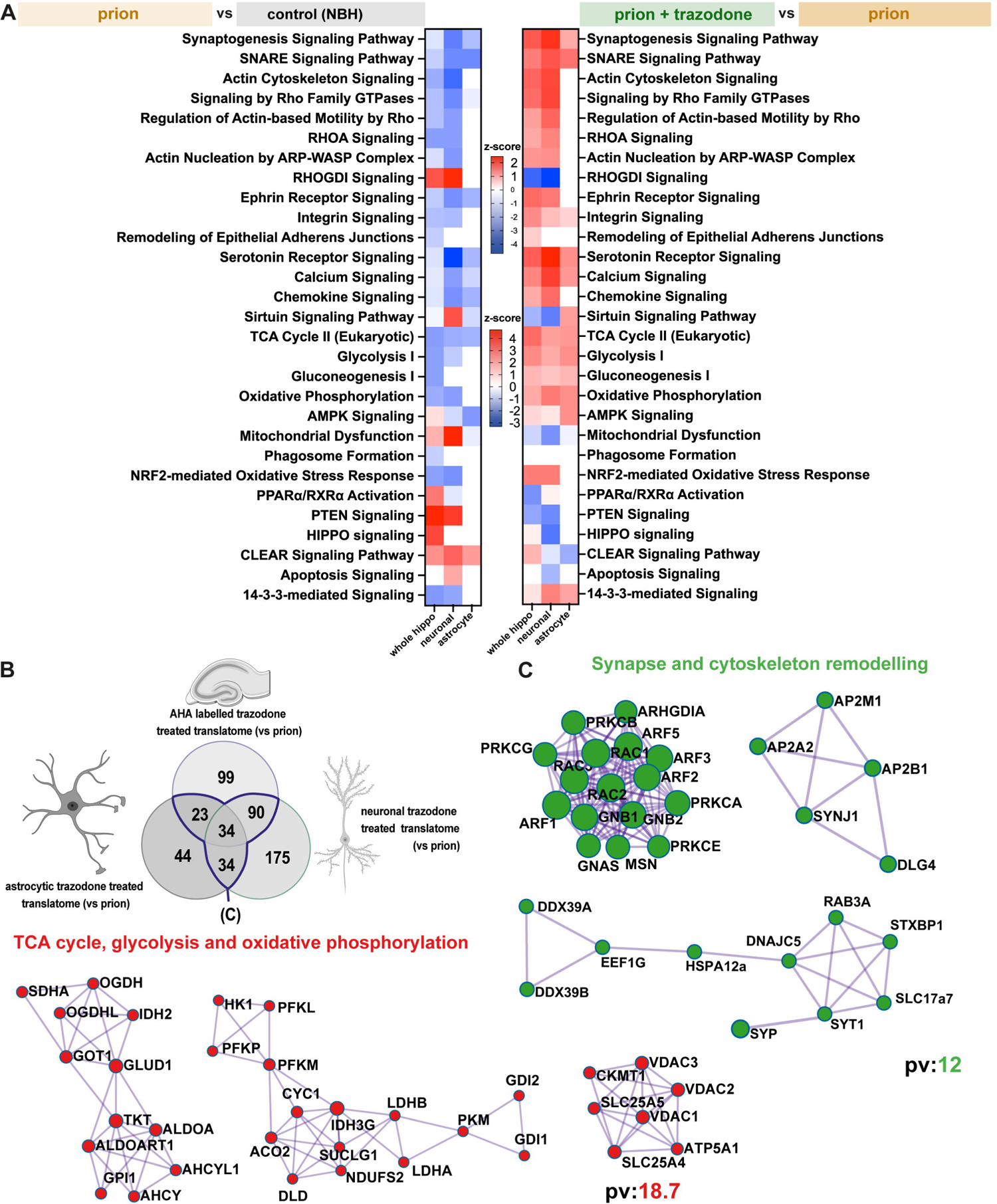
Synapse remodelling and mitochondrial function are the predominantly dysregulated pathways in prion disease and are restored by trazodone treatment. **(A)** heatmaps of significant pathways showing differences in z-score of prion (orange) vs control (NBH) (grey) translatomes (left), and heatmaps of significant pathways showing differences in z-score of prion+trazodone (green) vs prion (orange) translatomes (right). **(B)** Venn diagram of overlapping prion+trazodone translatomes from whole hippocampus, neurons and astrocytes, compared to prion alone (blue line highlights overlap). **(C)** Significant clusters of overlapping prion+trazodone translatomes, determined by metascape-string analysis, of significantly upregulated proteins compared to prion alone. The proteins clustered to synapse and cytoskeleton remodelling, and to the TCA cycle, glycolysis and OXPHOS. *p* value <0.05 for significant proteins; pv: -log_10_Pvalue.

Volcano plots comparing the translatomes for prion-disease versus controls for whole hippocampi, showed 141 proteins were significantly downregulated and 73 upregulated (**Fig. 3A**). For the CaMK2a-specific dataset, 229 proteins were downregulated and 31 were upregulated (**Fig. 3B**) and in the astrocyte-specific dataset, 110 proteins were downregulated and 18 were upregulated (**Fig. 3C**). In contrast, when comparing the nascent translatomes of prion+trazodone treatment to prion-only, the situation is reversed, with upregulation of 227, 230 and 123 proteins in whole hippocampi, neurons and astrocytes, respectively and downregulation of 20, 24 and 7, respectively (**Fig. 3A-C).** Altogether, the comparison of trazodone-treated compared to prion-diseased translatomes highlights recovery of the overall landscape of the diseased brain across different cell types. Consistent with this, the nascent translatomes of controls versus trazodone-treatment are relatively similar across the three groups with 45, 15 and 29 proteins upregulated and 80, 85 and 60 downregulated (**Fig. 3A-C**). The data are particularly compelling given the consistency between different mouse models, the tg37^+/-^ mice (FVB background, rapid disease course) and the NCAT::CaMK2a and NCAT::GFAP mice (C57/Bl6 background, slower disease course), and between whole hippocampal translatome from tg37^+/-^ mice with their mixed cell population and the neuron- and astrocyte-specific translatomes from the NCAT mice.

Proteins consistently downregulated in prion disease compared to controls include those related to synapse formation and maintenance, such as SYP or RAC1 and to ATP biosynthesis such as Pkm, Sdha, Vdac2 Slc25a4, in all samples (**Fig. 3A i, B i and C i**). Trazodone treatment upregulated (restored) many of these, including SYP, SYNGR1, DLG4 and SLC25A4, VDAC2, SDHA and PKM, and other mitochondrial proteins ATP5A1/J2, NDUFS1 or PC (**Fig. 3A ii, B ii, C ii and Supplementary table2**). Interestingly, some proteins including ATP-hydrolysis factors (ATP1B2, ATP8A1), amino acid importers (SLC1A4), cell adhesion proteins (CLDN11, VCL), mediators of autophagy (PIP4K2B) or protein ubiquitination (CAND1) were higher in trazodone-treated samples than controls, suggesting an overshoot or upregulation of these by trazodone treatment in disease (**Fig. 3A iii, B iii and C iii**). A few proteins upregulated in control compared to trazodone-treated whole hippocampi (UQCRC1, ATP1B2, MDH2 OR HSPA8) were downregulated in the cell-specific translatomes (HSPA8, ATP1B2 in neurons) and (MDH2, UQCRC1 in astrocytes).

Taken together, the data highlight the profound impact of prion disease on nascent translatome during early neurodegeneration seen at the level of both bulk hippocampal tissue as well as in hippocampal neurons and astrocytes, with profound impairment in synaptic and mitochondrial proteins, further defined below. Trazodone treatment restored many of these proteins, consistent with its neuroprotective effects.

### Trazodone restores dysregulated pathways controlling synapse and mitochondrial function

We further analysed significantly differentially expressed proteins represented in the volcano plots (**Fig. 3).** Using their log2-fold change values we applied ingenuity pathway analysis (IPA) to reveal up- or down-regulated canonical pathways for each of the nascent translatomes **(Fig. 4A).** Across all the prion versus control translatomes, many important pathways related to key aspects of hippocampal function were downregulated, including synaptogenesis signalling and other synapse-related pathways - SNARE signalling, actin cytoskeleton signalling and others. Another major group of downregulated pathways were related to mitochondrial function, including TCA cycle, glycolysis and oxidative phosphorylation (OXPHOS), although the latter two are not downregulated in the astrocyte-specific translatome. Upregulated pathways in prion-diseased animals versus controls include RHOGDI signalling - related to inhibition of actin Rho small GTPases; mitochondrial dysfunction and CLEAR signalling and in neurons specifically, apoptosis (**Fig. 4A**). These changes are reversed to varying degrees by trazodone treatment, both within the whole hippocampal translatomes of prion-diseased tg37^+/-^ mice, and within the neuron- and astrocyte-specific translatomes of prion-diseased NCAT mice. In particular, the data from prion+trazodone versus prion highlight the restoration of synaptogenesis, SNARE and actin cytoskeleton signalling pathways as well as Rho GTPases, RHOA, ephrin, integrin, serotonin signalling, the latter in whole brain and neurons, consistent with changes in these translatomes during disease. Trazodone treatment results in recovery also of mitochondrial pathways glycolysis, TCA cycle and oxidative phosphorylation signalling in all nascent translatomes, with reduction of mitochondrial dysfunction and neuronal apoptosis (see full list in **Supplementary Table 3)**.

We next looked at the specific proteins affected in all the trazodone-treated translatomes compared to vehicle-treated translatomes (**Fig. 4B; full details supplementary Table 4**) and analysed these by metascape-STRING^53^ (**Fig. 4C**). Consistent with the IPA analysis, this confirmed two main clusters of highly interacting proteins, one related to synapse and cytoskeleton remodelling and the other to mitochondrial biology: TCA cycle, glycolysis and OXPHOS (**Fig. 4C**).

### Quantitative and functional rescue of synapses and mitochondria by trazodone

For a morphological assessment of the proteomics results implicating synaptogenesis and mitochondrial pathway impairments, we used electron microscopy to assess synapse and mitochondrial numbers in the CA1 region in all groups of tg37^+/-^ (as per schematic in **Fig 1A**). Synaptic density was dramatically lowered in prion-disease (2.8 synapses/55µm^2^) compared to controls (6.5 synapses/55µm^2^; *p* < 0.0002), consistent with previous findings^21, 51, 58^. In contrast, two weeks of trazodone treatment significantly rescued synapse density (to 4.54 synapses/55µm^2^; *p =* 0.0096), although not to wild type levels (**Fig. 5A)**. Synaptic proteins, detected by immunoblotting, also increased (**Supplementary Fig. 4A**). In our previous work, we showed that trazodone restores memory^22^ in prion-diseased mice, a functional recovery consistent with its rescue of synapse number and protein levels demonstrated here.

**Figure 5.**
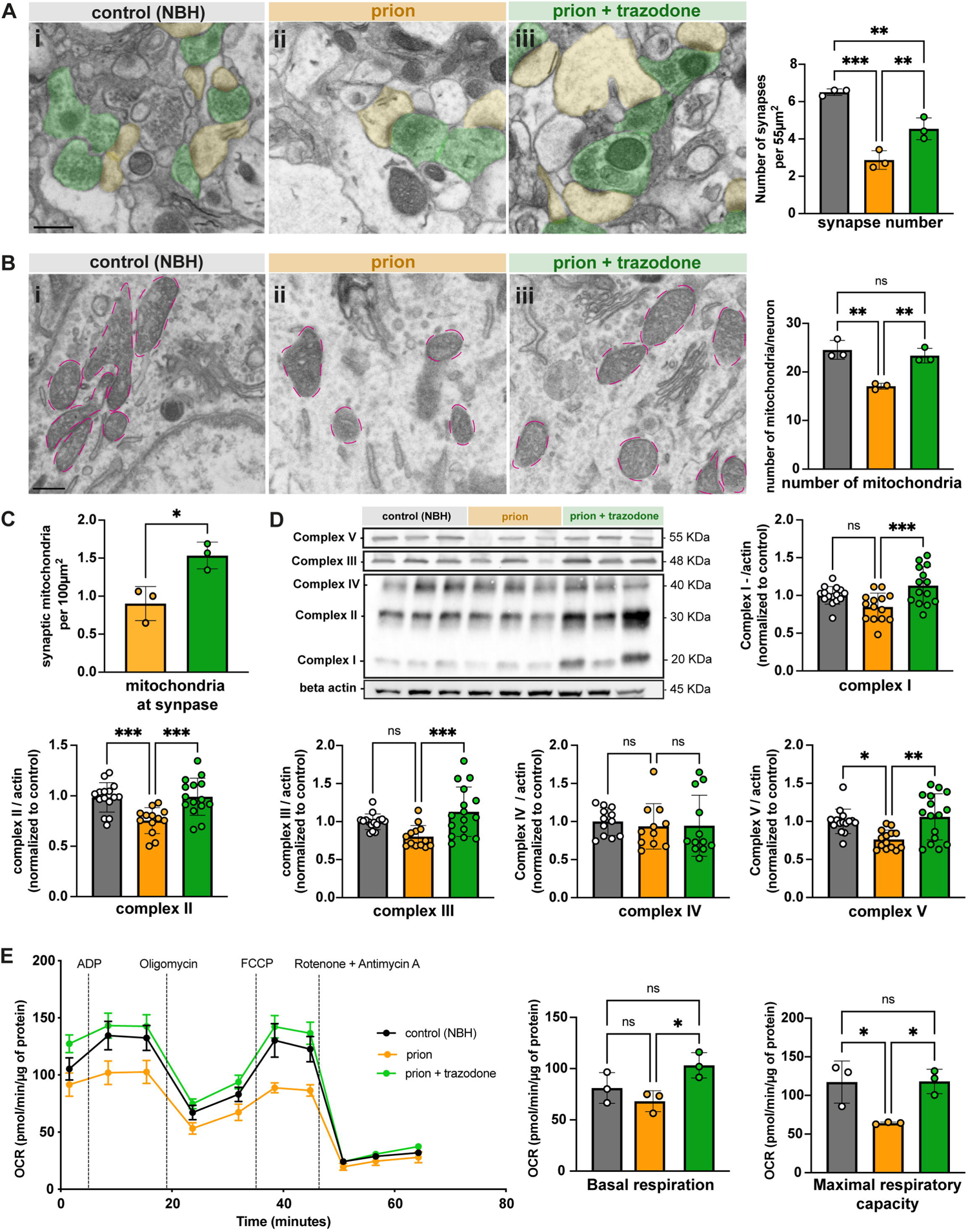
Trazodone treatment restores synapse and mitochondrial numbers and function in prion disease. **(A)** Representative SEM images from control (NBH) (**i**), prion (**ii**) and prion+trazodone mice (**iii**) from CA1 stratum radiatum, with dendrites pseudo-coloured in yellow; axons in green. Synapse number quantification of the hippocampal CA1 neuropil shows a decrease in prion disease (orange bars) and partial but significant restoration with trazodone treatment (green bars). **(B)** Representative SEM images from control (NBH) (**i**), prion (**ii**) and prion+trazodone (**iii**) from CA1 pyramidal neurons somata with mitochondria perimeter coloured in pink. Mitochondrial quantification in CA1 pyramidal neuron somas shows reduced number of mitochondria/neuron in prion disease (orange bars) and restoration to wild type levels with trazodone treatment (green bars). (**C**) Mitochondria counted within synapses expressed per 100µm^2^. **(D)** immunoblots and quantification (bar graph, right) of oxidative phosphorylation proteins (I,II,III, IV and V) showing restoration with trazodone treatment. (**E**) Mitochondrial stress test on isolated mitochondria from control (NBH), prion and prion+trazodone treatment groups at 10 w.p.i. as before (see Fig. 1A), showing oxygen consumption rates (OCR) from 0 to 65 minutes, with time points of stressor addition (ADP, oligomycin, FCCP and rotenone + antimycin A, respectively) labelled. Basal respiration was defined as the (initial) OCR value minus (rotenone + antimycin A) OCR value; maximal respiratory capacity was defined as the OCR value after FCCP injection minus (rotenone + antimycin A) OCR.. *p* values * < 0.05, ** < 0.001,*** < 0.0001. Scale bar A(i) and B(i): 1μm.

Mitochondrial numbers in the neuronal cell bodies were reduced in disease (17/pyramidal neuron) compared to controls (25/pyramidal neuron; *p =* 0.0018) and restored to wild type levels by two weeks trazodone treatment (23.3/pyramidal neuron; *p =* 0.0045) (**Fig. 5B**). Trazodone also restored mitochondria numbers within dendrites and axons of the CA1 neurons (**Fig. 5C**). In parallel, trazodone treatment produced significant recovery in oxidative phosphorylation protein complexes I, II, III and V, as detected by immunoblotting (**Fig. 5D**).

We next asked whether, independently of differences in mitochondria numbers, the lower levels of OXPHOS protein complexes in disease (seen both in the nascent translatomes of whole hippocampi, neurons and astrocytes (**Fig. 4A and 5B**) and in immunoblots (**Fig. 5D**) is reflected in a functional effect on mitochondrial respiration. Likewise, we asked if the quantitative changes induced by trazodone are reflected functionally. We performed mitochondrial stress testing Seahorse XF96 Analyzer (Agilent) on mitochondria isolated from whole hippocampi of tg37^+/-^ control (NBH), prion-diseased and prion+trazodone mice at 10 w.p.i., using equal numbers from each condition. Mitochondrial respiration, measured as oxygen consumption rate (OCR), was recorded at baseline and in response to different stressors. Basal mitochondrial respiration was not significantly different between controls (81.1pmol/min/µg) and prion-diseased samples (68.2pmol/min/µg, *p* = 0.47), but trazodone treatment resulted in a significantly higher OCR (103.1pmol/min/µg; *p =* 0.0342) (**Fig.5E**). No differences in respiration were observed between groups after injection of ADP or oligomycin, consistent with true OXPHOS impairment (**Supplementary Fig. 4C**). Injection of FCCP, an uncoupler that depolarizes mitochondria and maximises OCR, resulted in significantly lower OCR in prion mitochondria than in controls (64.1 vs 117.2 pmol/min/µg, *p* = 0.0274), which was restored to control levels by trazodone treatment (118 pmol/min/µg, *p* = 0.0257) (**Fig 5E**). Mitochondria from prion-diseased brains failed to increase their respiratory rate at all with FCCP, consistent with the reduction in OXPHOS proteins rendering ATP production maximal at baseline, with no spare capacity. When exposed to rotenone and antimycin A, which inhibit complexes I and III of the electron transport chain, OCR levels of all groups were drastically lowered below basal respiration with no differences between groups, as expected (**Fig 5E**). Together, the data show that prion disease affects mitochondrial biology through reduction in net numbers and protein levels, as well as functionally, with reduced respiration - independent of number. Trazodone treatment rescues these defects, restoring mitochondrial capacity to sustain neurons and likely other cells in the hippocampus with crucial ATP production. The nascent translatome changes at the synapse and in mitochondria and their reversal by trazodone are in **Fig 6**.

**Figure 6.**
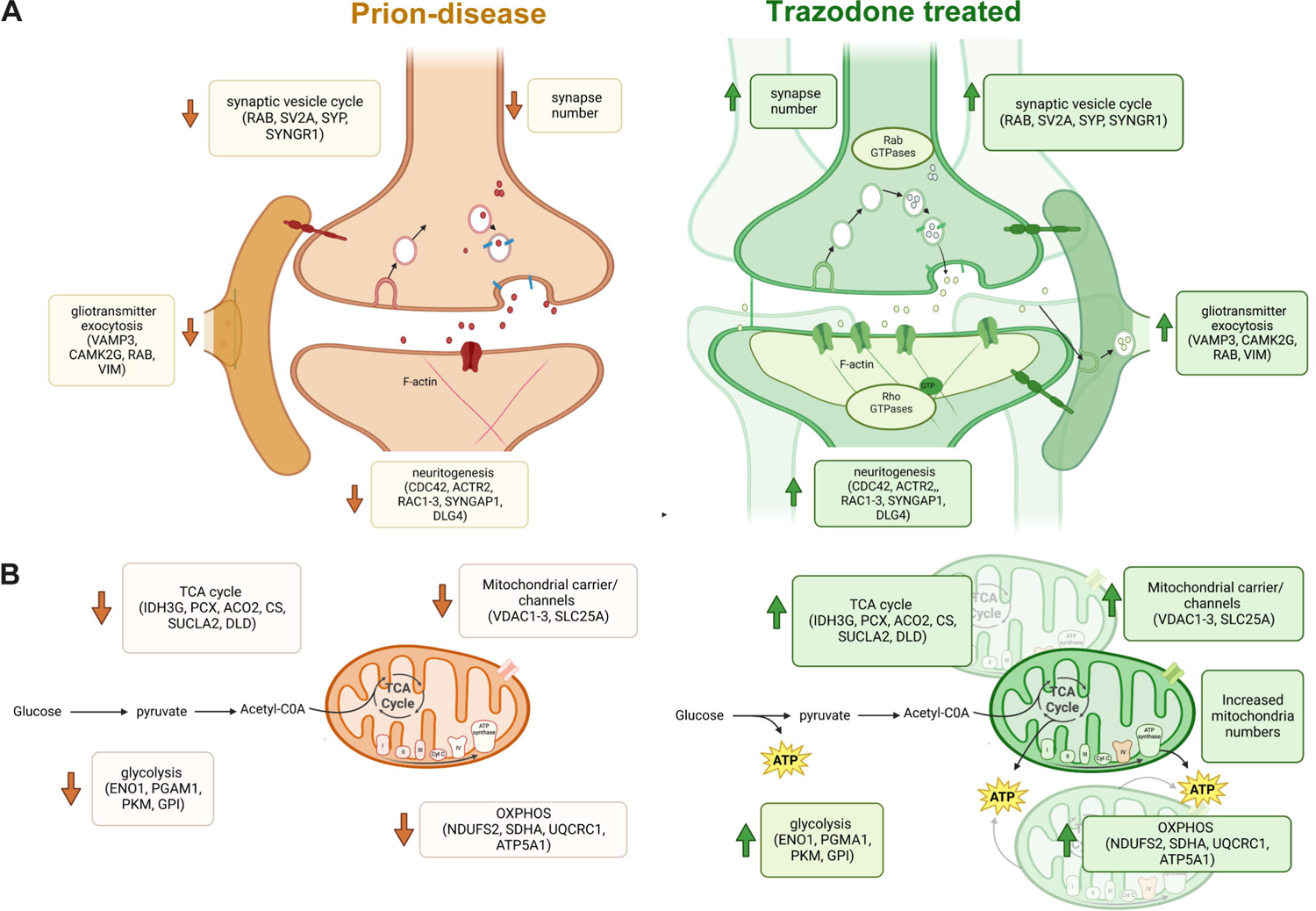
Trazodone treatment in prion disease. Schematic summarizing pathway and major proteins downregulated changes in the nascent proteome during prion disease and recovery after trazodone treatment for **(A)** synapses and **(B)** mitochondria.

## Discussion

Repression of protein synthesis rates in neurodegenerative disease is increasingly accepted to be a contributing factor to impaired memory formation, synapse loss and neuronal demise across the spectrum of these disorders. Several pathways contribute to the failure of protein homeostasis (proteostasis), including those controlling both the initiation^59^ and elongation^60^ phases of translation, ribosomal function^61–64^, and protein degradation^65^. Dysregulation of the control of the initiation phase of translation, in particular, via the PERK branch of the UPR occurs across the spectrum of human neurodegenerative disease^10^. In mouse models of neurodegenerative disorders, from Alzheimer’s, Parkinson’s, tauopathies, ALS and prion disease, over-activation of this pathway is associated with pathology and clinical signs. Genetic or pharmacological reversal restores global protein synthesis rates and is markedly neuroprotective despite high levels of ‘toxic’ misfolded proteins, highlighting the importance of helping restore proteostasis to mitigate the effects of proteotoxicity^11, 12, 14–25, 34, 48^. Restoring translation at the level of elongation is also protective in Alzheimer’s mouse models^35–37^, further speaking to the importance to neuronal survival of maintaining a healthy proteome. Yet the precise nature and identity of these dysregulated proteins has only recently begun to be explored^43–45^ and is an important focus for an increased understanding of the subcellular mechanisms and processes that lead to neuronal dysfunction and death.

In this study we wanted to define the altered nascent proteome during disease - prior to overt neurodegeneration - and its response to treatment. Our hypothesis was that if UPR activation, known to be associated with synaptic failure and neuronal demise, leads to dysregulation of a specific set of proteins, then UPR-targeted treatments should, at least in part, reverse this. We were interested in both the neuronal and astrocytic specific proteomes as both cell types are critically involved in synapse formation and function, and modulating UPR activation in each is equally neuroprotective^21, 24^. We found proteins involved in two major sets of biological processes and subcellular structures to be dominantly affected by the disease processes: those involving the synapse and the mitochondria.

Synaptic dysfunction is the main cause of memory-associated cognitive deficits in neurodegenerative disease and is a prequel to neuronal demise^66, 67^. Our data show that during prion disease, when synapse loss is established but neuronal loss is imminent - 10 w.p.i. in tg37^+/-^ mice and 18 w.p.i. in NCAT mice - that levels of predominantly neuronal synaptic proteins, both presynaptic (including SYN2, STX1B, SYNGR1, SYP) and postsynaptic (including SYNGAP1, DLG4, SLC6A11) are lowered by disease but restored by trazodone treatment (**Fig. 3 and Supplementary Table 2**). Concurrently, we observed an increase in levels of RAC1 and CDC42, Rho GTPases known for their role in F-actin assembly and disassembly important in bouton and dendritic spine formation^68, 69^ by trazodone (**Fig. 3 and Supplementary Table 2**), suggesting that synaptic remodeling is also restored. Presynaptic membranes are also affected with reduced levels of small Rab GTPases, which play a key role in the loading of neurotransmitters into synaptic vesicles^70, 71^, again reversed by trazodone treatment. Trazodone also reduced apoptotic signaling pathways upregulated in neurons during disease (**Fig. 4 and Supplementary table 4**). Astrocytes are also key for synapse maintenance. They closely interact with neurons at both the pre- and post-synapse, in a structure known as the tripartite synapse^72^. This is critical in development to establish neuronal connectivity, and in adult brains to integrate synaptic activity, form/eliminate synapses and to release gliotransmitters such as ATP, glutamate or serine in a Ca^2+^ dependent manner^72^. Changes to gliotransmitter release have been shown to have an impact on neuronal function and are believed to contribute to neurodegenerative disease^73^. Prion disease produces a UPR-reactive astrocytic state that leads to a non-cell autonomous neuronal degeneration in prion disease via an altered astrocytic secretome that is reversed by UPR inhibition with recovery of synaptic and neuronal numbers and extended survival^24^. Our BONCAT analysis of trazodone-treated astrocytes showed a restoration of key proteins involved in SNARE signalling (RAB2A, NSF, SYT3, VAMP3, ERC2), key for astrocytic exocytosis, supporting the hypothesis that trazodone is rescuing astrocyte neurotrophic support. Also restored are integrin signalling (ARF5, ARHGDIA, PFN2), important for tripartite synapse function (**Fig. 4 and Supplementary Table 2**). The data thus provide detailed mechanistic underpinning at multiple levels of synaptic function and structural plasticity in both neurons and astrocytes for the functional recovery in memory we and others have previously shown with trazodone in prion-diseased^22^ and tauopathy rTg4510 mice^22, 34^, as well as protection from neurodegeneration. The protection from synapse loss seen in SEM images (**Fig. 5a**) by trazodone treatment is also explained by these changes in the proteome.

Also essential for both synaptic and neuronal function are healthy mitochondria. Mitochondrial dynamics, mobility and turnover are perturbed, and TCA cycle proteins and OXPHOS complexes are downregulated in prion and other neurodegenerative diseases in humans and mouse models^74–82^. Mitochondria are commonly found at nerve terminals where they help maintain neurotransmission at the pre-synapse through Ca^2+^ efflux and the production of ATP^83, 84^. Both GTPase activity and increased protein synthesis rates in neurons require ATP and GTP, synthesized by mitochondria. The importance of glycolysis and mitochondrial metabolism for neurons and specifically for synapse function is well established^74–78, 83–85^. In a similar fashion, dendritic mitochondria are also essential at ER-mitochondrial contact sites, which regulate ATP and Ca^2+^ dynamics for the morphogenesis and plasticity of synaptic spines^83, 84^. Our BONCAT analyses and validation experiments reveal that trazodone treatment increases mitochondrial activity by increasing the expression of different proteins related to glycolysis (PKM, GPI, TPI1), TCA cycle enzymes (PCX, CS, DLAT, LDHA/B) as well as OXPHOS complexes (NDUFS1, SDHA, UQCRC2, MT-CO2, CYC1, ATP5J2) that are down-regulated in prion disease (**Fig. 3 and Supplementary table 2**), as also seen in the hamster tg7 prion model which shows downregulation of OXPHOS complex I and V, as well as proteins involved in TCA cycle^79^. Consistent with this, trazodone protects from mitochondrial loss with restoration of reduced numbers to wild type levels both within neuronal and within dendrites (**Fig. 5B,C**). Critically, these changes in protein levels are reflected in mitochondrial functional output, with lower maximal respiratory capacity in mitochondria from diseased brains compared to controls (as also seen in tg7 prion hamsters^79^), which trazodone was able to reverse completely, restoring normal mitochondrial respiratory function as well as numbers (**Fig. 5E**).

Astrocytes are an important hub for metabolism in the brain, providing many key metabolites to support synaptic and neuronal function, including ATP^80^. Trazodone restored astrocytic mitochondrial proteins including those of TCA cycle and glycolysis (PKM, TPI1, IDH3G, SUCLG1) and oxidative phosphorylation (UQCRC1, MT-CO2, ATP5B, CYC1). We hypothesize that this restoration of key metabolic processes in astrocytes aids in the synaptic protection discussed above. Importantly, mitochondrial dysfunction, ultimately leading to a lowering in ATP production, is not exclusive to prion disease but has been described throughout the spectrum of neurodegenerative diseases^76–79, 81, 82, 86, 87^, including Alzheimer’s^44, 45, 76, 77, 82, 86^, Parkinson’s^88^, tauopathies^43, 87^ and ALS^78^.

The proteome in various neurodegenerative diseases has been previously examined, including in prion disease, notably in human brains, where downregulation of pathways associated with Parkinson’s, Alzheimer’s disease, lysosomes and OXPHOS are reported^89^. Mouse prion bulk proteomic data from whole brains showed changes in calcium signalling^41^, and neuroinflammation^42^ in terminal disease, but these studies don’t address the acute translational changes, which require analysis of the nascent translatome and are also not cell-specific. More recently and relevant to our study, the *de novo* translatome has been examined using NCAT in mouse models of frontotemporal dementia (K3 FTD)^43^ and APP/PS1 model of Alzheimer’s disease^44, 45^. In K3 mice mice^43^, AHA labelling of the whole brain proteome over 16 hours in 5 months old mice showed, similar to our findings, a large reduction in neuronal protein synthesis rates measured by FUNCAT ^43^. Proteomics analysis showed reduced differentially expressed proteins related to OXPHOS complexes, the TCA cycle, microtubules and actin cytoskeleton, synaptic proteins and ribosomal proteins, confirmed in rTg4510 mice, again similar to our findings and to results from hippocampal slices from APP/PS1 mice^44^, examined by AHA labelling over 5 hours combined with SILAC (BONLAC) which again revealed changes in synapses, proteasome, lysosome, OXPHOS and ribosomal proteins^44^. A separate study in APP/PS1 mice, heavy labeled AHA hippocampal proteome showed similar changes^45^. Importantly, independent of neurodegenerative disease model used, anatomical area analysed and approach used to label the nascent translatome, there is a consistent and comprehensive overlap of proteins and cellular functions dysregulated across prion, tauopathy and Alzheimer’s disease^43–45^, stressing the mechanistic commonality of fundamental neurodegenerative processes across these disorders. However, none of these studies examined the response of the nascent translatome to therapies directed at restoring translation. Here, we have gained a deeper understanding of the proteins dysregulated by changing protein synthesis rates and their subsequent rescue with trazodone in both neurons and astrocytes in prion disease, supporting trazodone’s neuroprotective effect is due to a synergic protection in multiple cell types in the brain.

In conclusion, the data bring new insights into how translational repression during disease is so devastating to synaptic function and neuronal survival, via the chronic depletion of essential synaptic and mitochondrial proteins and ATP, resulting in loss of synapses, mitochondria and ultimately, neurons. They also bring new understanding of the specific mechanism of action of trazodone in mediating neuroprotection, through restoration of these key proteins essential for neural function and resilience. Given the marked commonalities we find with the dysregulated proteomes in Alzheimer’s and tauopathy models, and the known underpinning of UPR dysregulation across these disorders, our results on treatment with trazodone have wide implications for therapy of neurodegenerative disorders.

## Data availability

Processed data is available at **Supplementary Table 2-4,** Raw data is available via ProteomeXchange consortium with identifier PXD042577.

## Acknowledgements

We thank <u>Erin Schuman</u> of MPI Frankfurt for the NCAT mice and assistance with optimisation of BONCAT technology.

## Supplementary material

Supplementary Figure 1 - FUNCAT analysis of NCAT::CamK2Cre and NCAT::GFAPCre hippocampi

Supplementary Figure 2 - BONCAT elutions - quality controls

Supplementary Figure 3 - Hippocampal nascent translatomes - quality control and PCA charts

Supplementary Figure 4 - Trazodone restores synaptic proteins and mitochondria in prion disease.

Supplementary Table 1 - MaxQuant parameters

Supplementary Table 2 - Differential expression analyses

Supplementary Table 3 - IPA analyses

Supplementary Table 4 - proteins upregulated in trazodone treatment across translatomes compared to prion disease.

---

**Supplementary figure 1.** FUNCAT analysis. **(A)** Schematic of NCAT::GFAP/CaMK2a transgenic mice. **(B)** Schematic showing progression of prion disease, ANL labelling and trazodone treatment in NCAT::CaMK2a and NCAT::GFAP mice. **(C)** Representative images of FUNCAT analysis of the CA1 region from NCAT::CaMK2a (**i-iii**) and NCAT::GFAP (**iv-vi**) mice. **(D)** Representative FUNCAT images from CA1 of AHA labelled control **(i-iii)** and negative AHA clicked animal **(iv-vi)** showing no FUNCAT signal in negative controls.

**Supplementary figure 2.** BONCAT elutions. **(A)** Representative elutions of the whole hippocampal, neuronal and astrocytic nascent translatomes, compared to negative controls. **(B)** ANL log2 intensity of detected proteins compared to negative AHA/ANL controls. **(C)** Heatmap of cell-specific LFQ scaled intensities markers identified on hippocampal, neuronal and astrocytic translatomes**. (**C): control (NBH), P: prion and T: prion+trazodone.

**Supplementary figure 3.** Hippocampal nascent translatome. **(A)** Schematic of click chemistry performed on AHA/ANL containing proteins analysed through either FUNCAT or BONCAT. **(B)** Immunoblot illustrating NCAT::GFAP ANL incorporation in control (NBH) (black line), prion (orange line) and prion + trazodone (green line) mice. **(C)** Immunoblot quantifications of AHA and ANL incorporations for whole, neuronal and astrocytic hippocampus nascent translatomes, normalized to negative control. **(D)** Principal component analysis plots of whole, neuronal and astrocytic hippocampus from all nascent translatomes. *P* values: * < 0.05, ** <0.001, *** <0.0001,**** <0.00001.

**Supplementary figure 4.** Trazodone restores synaptic proteins and mitochondria in prion disease. **(A)** Immunoblots and quantification of synaptic proteins (SYP and VAMP2) in 10w.p.i tg37^+/-^ control, prion and prion+trazodone mice. **(B)** Average mitochondrial perimeter, area and frequency perimeter distribution in CA1 pyramidal neurons. (**C**) OCR from control (NBH), prion and prion+trazodone after ADP injection (left) and after oligomycin injection (right). (**D**) Images of isolated mitochondria from (i) control, (ii) prion and (iii) prion+trazodone tg37^+/-^ mice using mitotracker dye. *p* values: * < 0.05, ** <0.001, *** <0.001.

