## Supplementary figures and images for "Trazodone rescues dysregulated synaptic and mitochondrial nascent translatomes in prion neurodegeneration"

### Supplementary Fig.1

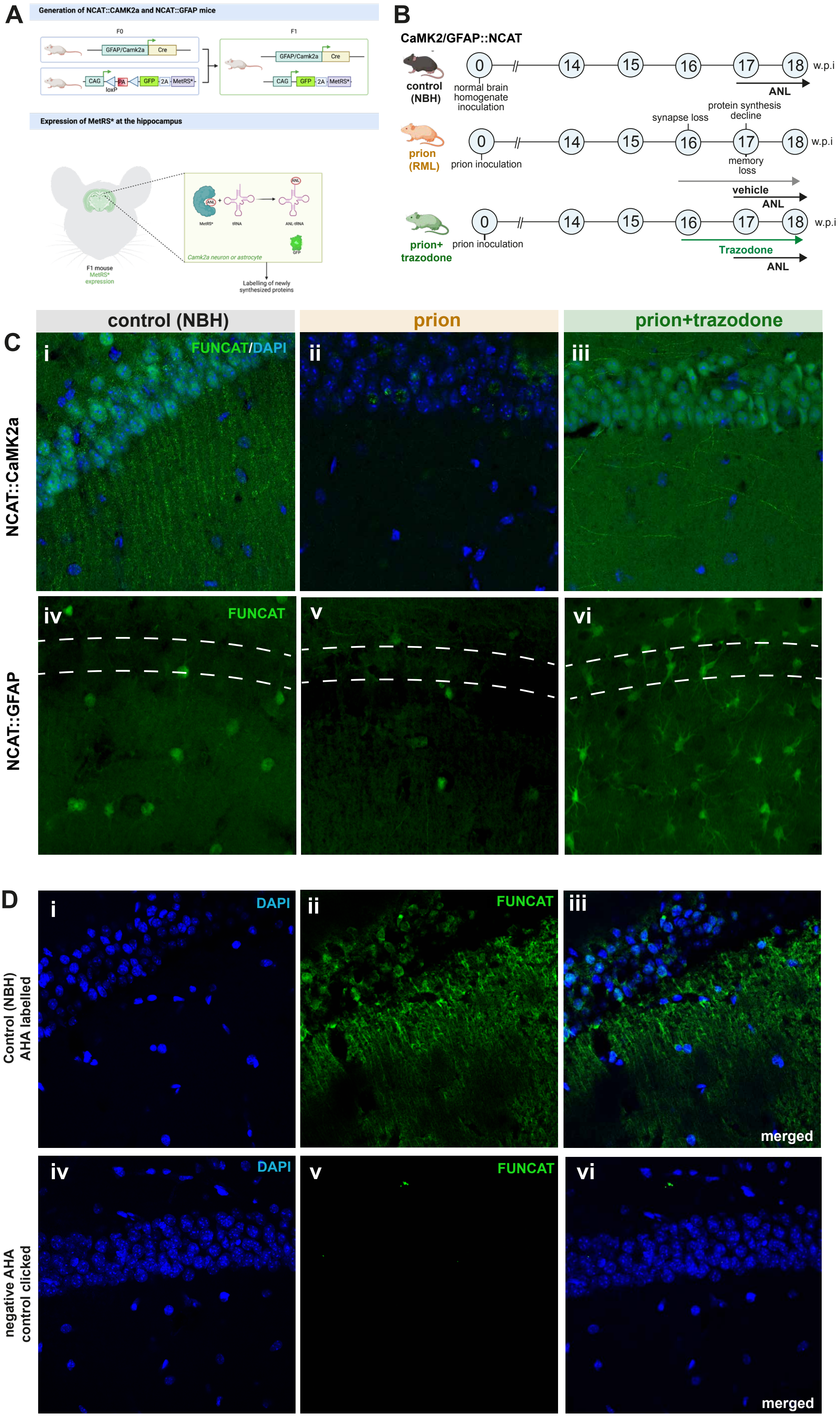

### Supplementary Fig.2

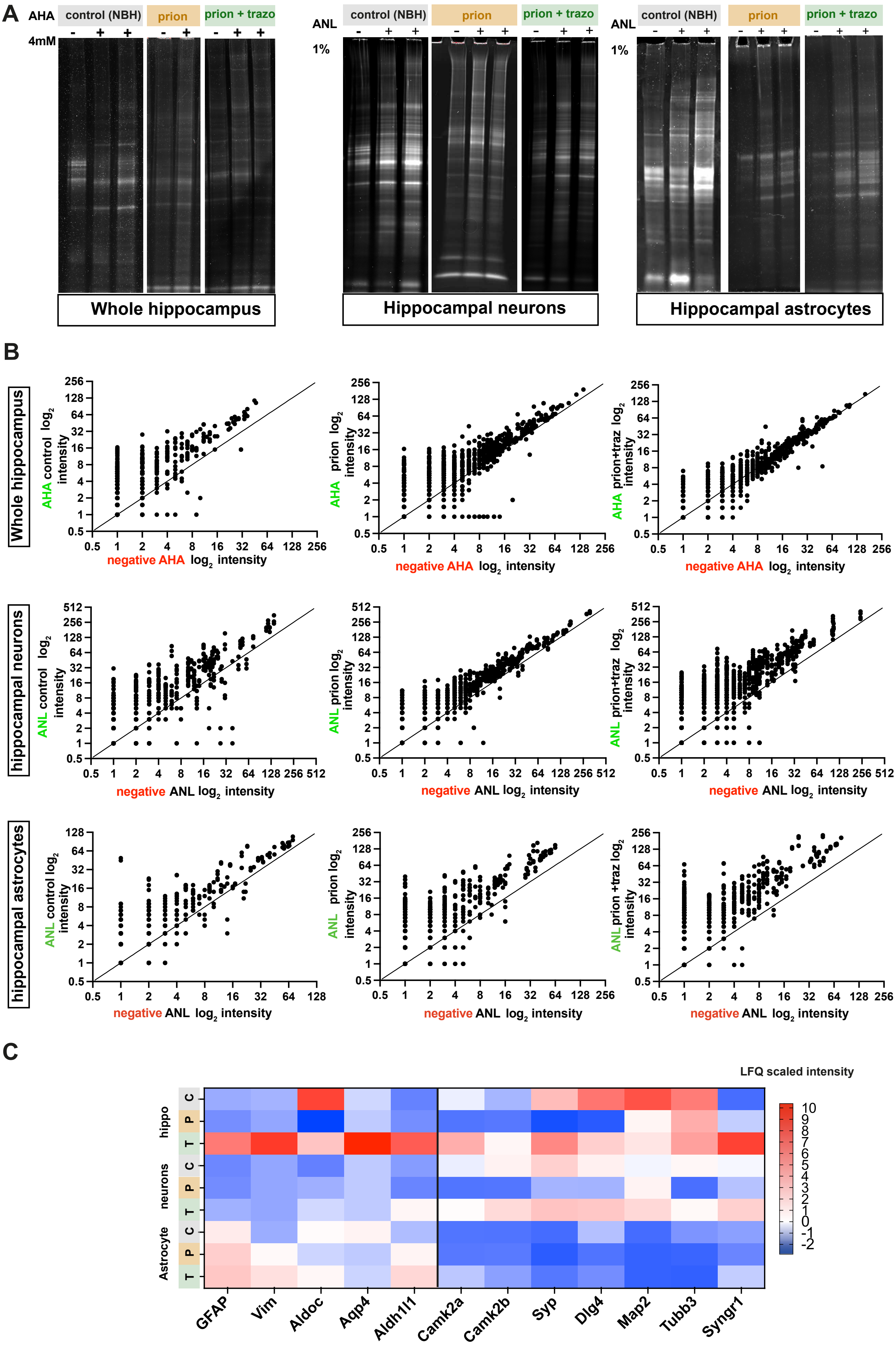

### Supplementary Fig.3

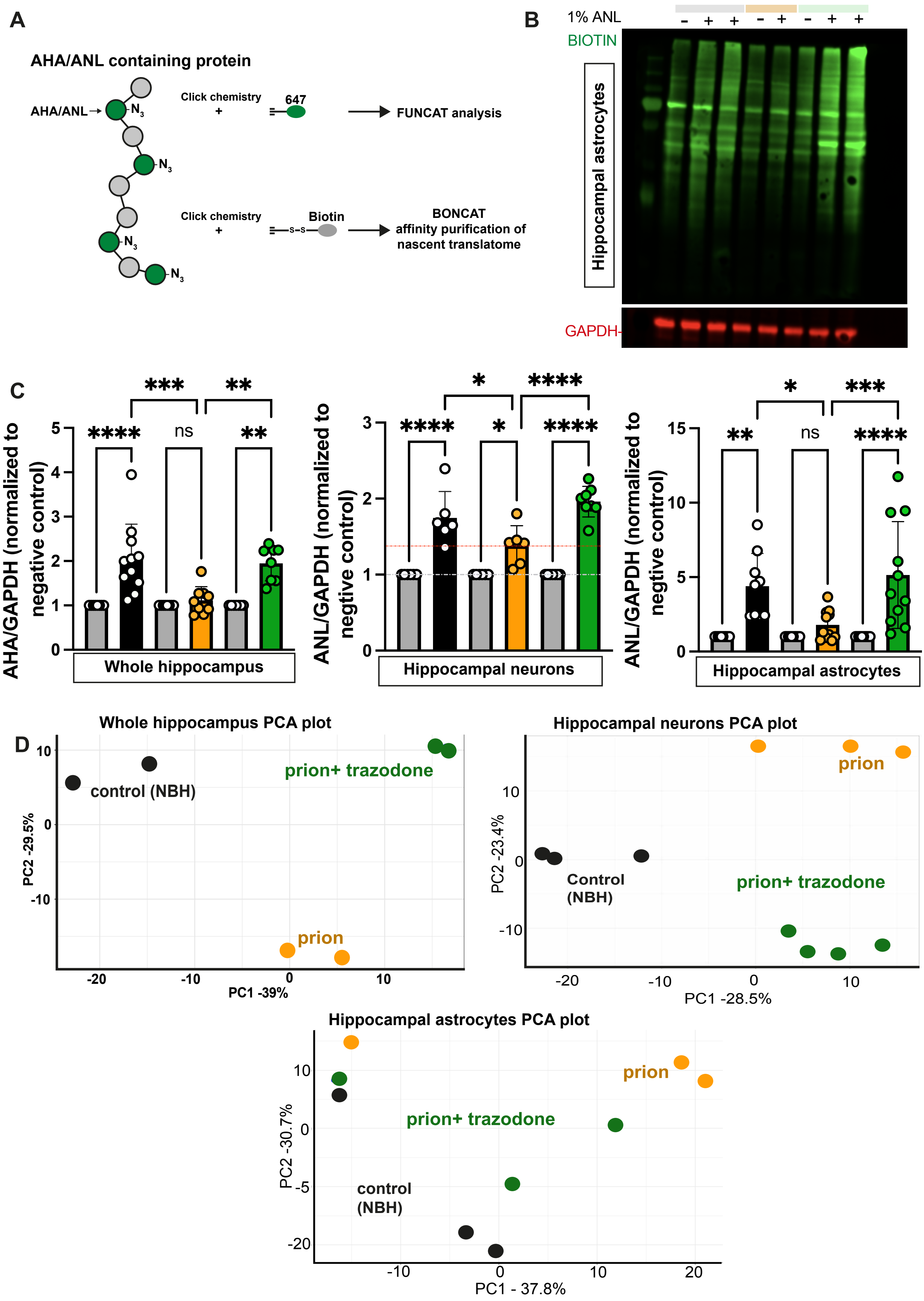

### Supplementary Fig.4

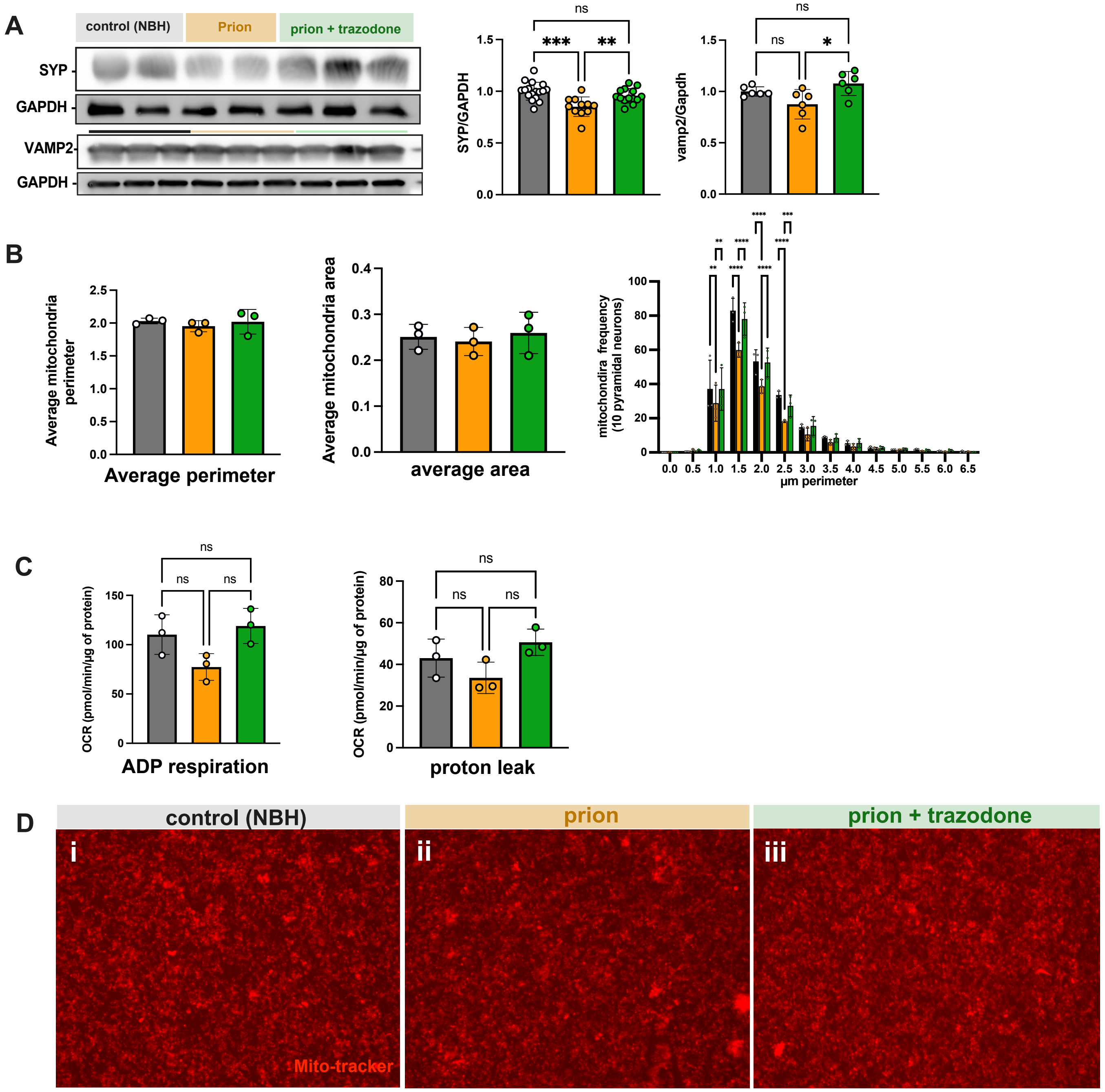
